# Glucagon Receptor-mediated Regulation of Gluconeogenic Gene Transcription is Endocytosis-dependent in Primary Hepatocytes

**DOI:** 10.1101/2021.09.03.458925

**Authors:** Jan Mikhale B. Cajulao, Mark E. von Zastrow, Erica L. Sanchez

**Affiliations:** San Francisco State University, Department of Biology; University of California San Francisco, Department of Psychiatry

## Abstract

A number of G protein-coupled receptors (GPCRs) are now thought to use endocytosis to promote cellular cAMP signaling that drives downstream transcription of cAMP-dependent genes. We tested if this is true for the Glucagon Receptor (GCGR), which mediates physiological regulation of hepatic glucose metabolism via cAMP signaling. We show that epitope-tagged GCGRs undergo clathrin and dynamin-dependent endocytosis in HEK293 cells after activation by glucagon, and transit via EEA1-marked endosomes shown previously to be sites of GPCR/Gs-stimulated production of cAMP. We further show that endocytosis potentiates cytoplasmic cAMP elevation produced by GCGR activation and promotes transcription of *PCK1*, the gene which encodes the enzyme catalyzing the rate-limiting step in gluconeogenesis. We verify endocytosis-dependent induction of *PCK1* expression by endogenous GCGRs in primary hepatocytes, and show similar control of two other gluconeogenic genes (*PGC1α* and *G6PC*). Together, these results implicate the endosomal signaling paradigm in metabolic regulation by glucagon.

## Introduction

G protein-coupled receptors (GPCRs) comprise the largest family of membrane proteins. GPCRs activate heterotrimeric G proteins in response to ligand-induced activation, then typically undergo regulated phosphorylation and bind to beta-arrestins. These events both attenuate GPCR-mediated activation of G proteins and promote receptor internalization. According to the classical view, GPCR signaling is restricted to the plasma membrane and internalized receptors are functionally inert (Harden et al., 1980). An emerging revision to this classical paradigm is that GPCRs have the potential to also signal after endocytosis (Calebiro et al., 2009; Irannejad et al., 2013; Pavlos & Friedman, 2017; Stoeber et al., 2018). Several GPCRs that initiate cAMP signaling through Gs (Rosenbaum et al., 2009) have been shown to couple to Gs both from the plasma membrane and in endosomes. These Gs-coupled GPCRs use activation from endosomes to promote downstream signaling to the nucleus for control of cAMP-dependent gene expression (Irannejad et al., 2013; Peng et al., 2021; Tsvetanova & von Zastrow, 2014). The full physiological and biological effects of this second wave of cAMP generation are an area of active investigation.

Endocytosis has been shown to promote cAMP-dependent signaling to the nucleus and transcriptional control by Gs-coupled polypeptide hormone receptors (Godbole et al., 2017; Tsvetanova & von Zastrow, 2014). For example, the Thyroid Stimulating Hormone Receptor must internalize for maximum transcription of genes that are part of the physiological response to thyroid stimulating hormone (Godbole et al., 2017). In developing oocytes, the Luteinizing Hormone Receptor internalizes to induce a second wave of cAMP signaling to the nucleus that promotes resumption of meiosis before ovulation (Lyga et al., 2016). The degree to which endocytosis-dependent downstream signaling is initiated remains unknown for other Gs-coupled polypeptide hormone receptors, such as the Glucagon Receptor (GCGR).

This study set out to explore whether endocytosis of GCGR was required for the transcriptional regulation of gluconeogenic gene expression. Blood glucose homeostasis is largely regulated by hormones, including insulin and glucagon, which have opposing physiological effects (Janah et al., 2019). While insulin’s effect is to reduce blood glucose during hyperglycemia, glucagon works to raise blood glucose levels during hypoglycemia (Janah et al., 2019). Glucagon is a peptide hormone released by the alpha cells of the pancreas that acts on GCGR expressed in the liver to stimulate gluconeogenesis and raise blood glucose content (Janah et al., 2019). A previous *in vivo* study found that a five day siRNA inhibition of endocytosis in mouse livers resulted in a loss in hypoglycemic homeostasis and a reduction of gluconeogenic gene transcription (Zeigerer et al., 2015). However, the role of GCGR endocytosis was unexplored. Therefore, GCGR is a GPCR whose subcellular signaling mechanisms have not yet been fully elucidated. Understanding the mechanisms and nuances of GCGR signaling could yield valuable insight for the receptor’s pharmacology and treatment of various metabolism-related diseases (Campbell & Drucker, 2015; Janah et al., 2019).

We hypothesized that the endocytosis of the glucagon receptor is required for transcriptional regulation of key gluconeogenic target genes. This study confirms that GCGR localizes to early endosomes upon stimulation and also demonstrates that endocytosis is required to elicit maximal cAMP production. Moreover, GCGR endocytosis is required for maximal GCGR-dependent transcription of gluconeogenic genes in primary hepatocytes. Our results reveal a new and previously unappreciated understanding of GCGR subcellular signaling.

## Materials and Methods

### Cell culture, expression constructs, and transfections

Human Embryonic Kidney 293 (HEK293) cells (ATCC CRL-1573) were cultured in complete growth Dulbecco’s modified Eagle’s medium (DMEM, Gibco), supplemented with 10% fetal bovine serum (UCSF Cell Culture Facility). Cells were passaged using PBS-EDTA and maintained at 37°C and 5% CO_2_. Transfections were performed using Lipofectamine 2000 (Life Technologies) according to the manufacturer’s protocol. Cells were transfected 48 hours before experiments. Primary cultures of mouse hepatocytes from male C57BL/6J mice were a generous gift from the UCSF Liver Center. Immediately following isolation, hepatocytes were plated on collagen coated cell culture plates and grown for no longer than 48 hours for all experiments (Corning BioCoat Collagen I, Cellware 6 well plate 356400). Hepatocytes were grown in DMEM supplemented with 10% fetal bovine serum and 1% Penicillin-Streptomycin (UCSF Cell Culture Facility). Microscopy experiments utilized the pCMV6-GCGR-MycDDK expression construct (OriGene Technologies, MR207767). All subsequent experiments used the generated signal sequence Flag-tagged glucagon receptor, pcDNA3.1(+)_SSF-GCGR, expression construct. Stable cell lines were generated and maintained by drug selection with Geneticin® Selective Antibiotic, G418 (Life Technologies, Cat# 10131035).

### Drug Treatment and siRNA

The highly potent dynamin inhibitor, Dyngo-4A (Abcam, ab120689), was added 15 minutes prior to agonist treatment, at a final concentration of 30 μM, from a 30 mM stock dissolved in DMSO. Vehicle control was prepared by adding DMSO without Dyngo-4A. For siRNA transfections, cells were seeded at 50% confluency in a 10 cm dish and transfected with 200 pmol siRNA (Qiagen, Germantown, MD) using RNAiMax Lipofectamine (Life Technologies, Invitrogen) according to the manufacturer’s protocol. Media was changed 24 hours post siRNA transfection and cells recovered for an additional 24 hours before experimentation. For siRNA control samples, cells were transfected with All Star Negative (siScramble) SI03650318 and for knockdown of Clathrin Heavy Chain, (siCHC17) 5’-AAGCAATGAGCTGTTTGAAGA-3’.

### Antibodies

Antibodies used were rabbit anti-Flag (Sigma), mouse anti-Flag M1 (Sigma), and mouse anti-EEA1 (BD Biosciences). Mouse anti-Flag M1 was conjugated to Alexafluor 647 for use in flow cytometry.

### Fixed cell confocal imaging

Cells for experiments were either transiently transfected with indicated construct(s) and processed 48 hours later, or were stable expression cell lines, selected with G418. Cells seeded on glass coverslips (Fisher, 12-545-100) in 12-well plates, were treated with or without Dyngo-4a followed by treatment with or without glucagon. After ligand treatment, cells were: (1) rinsed with PBS, (2) fixed by incubation in 3.7% formaldehyde (Fisher Scientific) diluted in PBS buffer for 20 minutes at room temperature, (3) blocked in 2% Bovine Serum Albumin (Sigma) in PBS with permeabilization by 0.2% Triton X-100 (Sigma), (4) labeled by the addition of primary antibodies diluted in blocking/permeabilization buffer for 1 hour, (5) secondary labeling was performed by addition of the following antibodies diluted in blocking/ permeabilization buffer for 30 minutes at room temperature: Alexa Fluor 647 or 488 donkey anti-mouse (Invitrogen), Alexa Fluor 647 or 488 donkey anti-rabbit (Invitrogen). Specimens were mounted using ProLong Gold antifade reagent (Life Technologies).

### Microscope image acquisition and image analysis

Fixed cells were imaged by spinning disk confocal microscope (Nikon Ti Inverted microscope with Yokogawa confocal scanner unit CSU22) using a 60X objective. Fluorescence intensities were quantified and analyzed using the computer program ImageJ (https://imagej.nih.gov/ij/). Images were saved as uncompressed 12-bit TIFF images, then loaded into ImageJ. Line scan analysis measured intensities along a manually drawn line across a cell, plotted using the Plot Profile function on ImageJ. Data were exported to Microsoft Excel and plotted on line graphs. Lines were drawn such that they intercepted endosomes and ran across the length of the cell. To quantify endocytosis of the Glucagon Receptor, colocalization was analyzed using the Coloc 2 ImageJ package. Cell outlines were manually drawn, and Pearson’s correlation coefficient (PCC) between indicated channels was calculated using Coloc 2. Mean PCC between samples were calculated, and tested for statistical significance using a twotailed unpaired *t* test (GraphPad Prism).

### Flow Cytometry

HEK293 cells stably expressing pcDNA3.1(+)_SSF-GCGR were treated with or without 1 μM Glucagon for 30 minutes at 37°C. Cells were then washed with PBS, and fixed with 4% formaldehyde. Cells were incubated with AlexaFluor647-conjugated (Invitrogen, A20173) M1 mouse anti Flag monoclonal (Sigma, F-3040) antibody at 4°C for one hour on a shaker. Cells were mechanically lifted and mean fluorescence intensity of 10,000 cells was measured by flow cytometry for each sample on a FACSCalibur (BD Biosciences). Internalization was calculated as the mean fluorescence intensity normalized to control samples. Each condition had three technical replicates per biological replicate.

### cAMP luminescence assay

HEK293 stably expressing pcDNA3.1(+)_SSF-GCGR cells transfected with GloSensor20F (Promega) were lifted and resuspended in imaging media (DMEM without phenol red supplemented with 30 mM HEPES; Gibco, 31053 and 15640 respectively). Cells were then incubated with D-luciferin (Gold Biotechnology St. Louis, MO; LUCNA-1g) at 37°C and 5% CO_2_ for one hour in 24 well plate (200 μL/well). In the last 15 min of incubation, samples were treated with or without Dyngo-4a. Immediately before imaging, cells were treated with 200 μL imaging media only or imaging media with Glucagon (1 μM) and placed into a 37°C heated light-proof chamber. Plates were then imaged every 10 seconds for 20 minutes. Images were acquired using a 512 × 512 pixel electron multiplying CCD camera (Hamamatsu Photonics, Japan; C9100-13) using μManager 1.4. Analysis was completed using the multiple ROI analysis tool in the Fiji implementation of ImageJ. ROIs were drawn around each well, and corresponding background ROIs were placed in an area without cells. Intensity over time was measured and corrected for background luminescence for firefly luciferase values. A ratio of firefly to background luminescence was calculated per well over time. The max (average) of the top 3 proximal values of each treated condition was determined, and each condition was normalized to the max of the control sample.

### RNA Extraction and Quantitative Reverse Transcriptase PCR

RNA was isolated from HEK293 cells using the QIAshredder and RNeasy Mini kit (Qiagen, 79654 and 74104, respectively). Briefly, after treatment, cells in a 6 well plate were placed on ice and washed once with cold PBS. Then cells were lysed, and RNA was extracted according to the manufacturer’s protocol. RNA was eluted in RNAse free water, and the concentration of each sample was determined. cDNA was generated from RNA using iScript cDNA Synthesis Kit (BioRad,1708891). Briefly, 1 μg of RNA was used per reaction primed with oligo dT. The reverse transcription reaction was performed according to the manufacturer’s protocol.

qRT-PCR was performed using a StepOnePlus (Applied Biosystems, Foster City, CA) instrument. cDNA generated from extracted RNA was used as the input for the qRT-PCR, with amplification by Power SYBR Green PCR master mix (Applied Biosystems, 4367659). Transcript levels were normalized to GAPDH or HPRT. The primer pairs are listed in table below.

#### qRT-PCR primers

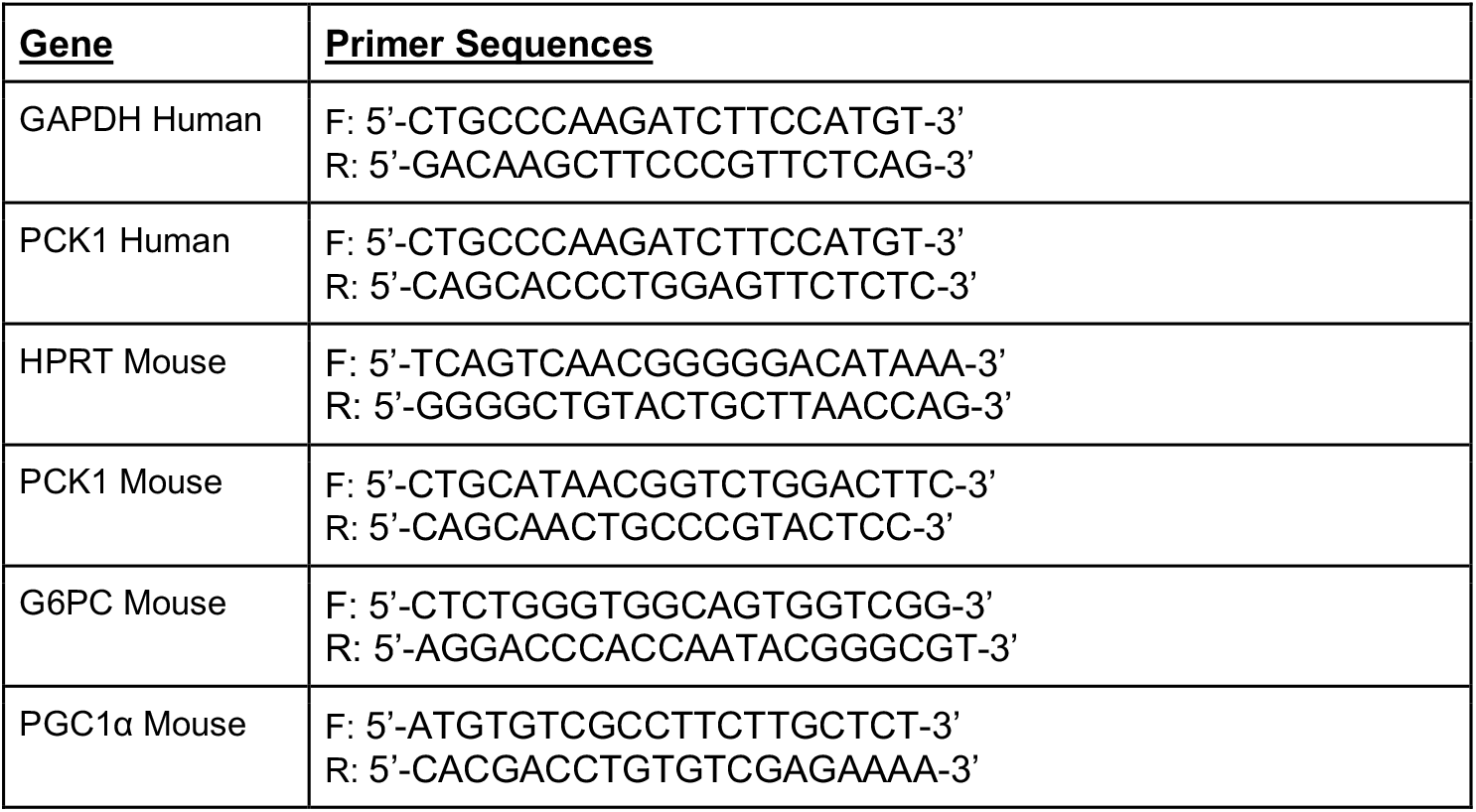

### Statistical analysis and reproducibility

All data are shown as mean ± SD or S.E.M. from at least three biologically independent experiments, unless otherwise indicated. Images are representative of at least three biologically independent experiments. Statistical analyses to determine significance were performed using Prism v.8 (GraphPad) for unpaired *t*-test, one-way analysis of variance (ANOVA).

## Results

### Stimulated Glucagon Receptor (GCGR) translocates to endosomes

Several G protein-coupled receptors signal at the cell surface, as well as within various intracellular locations after agonist stimulation. We hypothesized that upon stimulation with the glucagon peptide, that GCGR also traffics to the endosome. We first interrogated the trafficking and intracellular localization of GCGR after agonist stimulation. For these experiments, HEK293 cells were transiently transfected with the GCGR-MycDDK expression plasmid. GCGR and endosomal localization were visualized using antibody staining and confocal microscopy. Confocal images show GCGR primarily localizes to the plasma membrane in unstimulated cells (Figure 1A & B). Quantification of these images by line scan analysis show intensity peaks at the ends of the lines that correspond to the cell surface (Figure 1A & B, yellow arrowheads). When stimulated with glucagon, confocal images show that cytoplasmic GCGR increases, and accumulates in early endosomes. Line scan analysis showed that cytoplasmic GCGR intensities increase and create noticeable peaks that correlate with peak intensities of the endosomal marker EEA1 (Figure 1A & B, black arrowheads). To confirm that GCGR internalizes via the endocytosis mechanism, we first treated cells with Dyngo, a drug inhibitor that targets Dynamin (Figure 1A). We also performed these experiments upon siRNA knockdown of clathrin heavy chain (CHC17), a gene critical for endocytosis (Figure 1B). Upon treatment with glucagon, cells pre-treated with Dyngo failed to accumulate GCGR at endosomes. Similarly, GCGR did not accumulate at endosomes in cells where clathrin was knocked down. These are highlighted by the lack of intracellular GCGR puncta, and intensities that correlate with EEA1 (Figure 1B).

**Figure 1:**
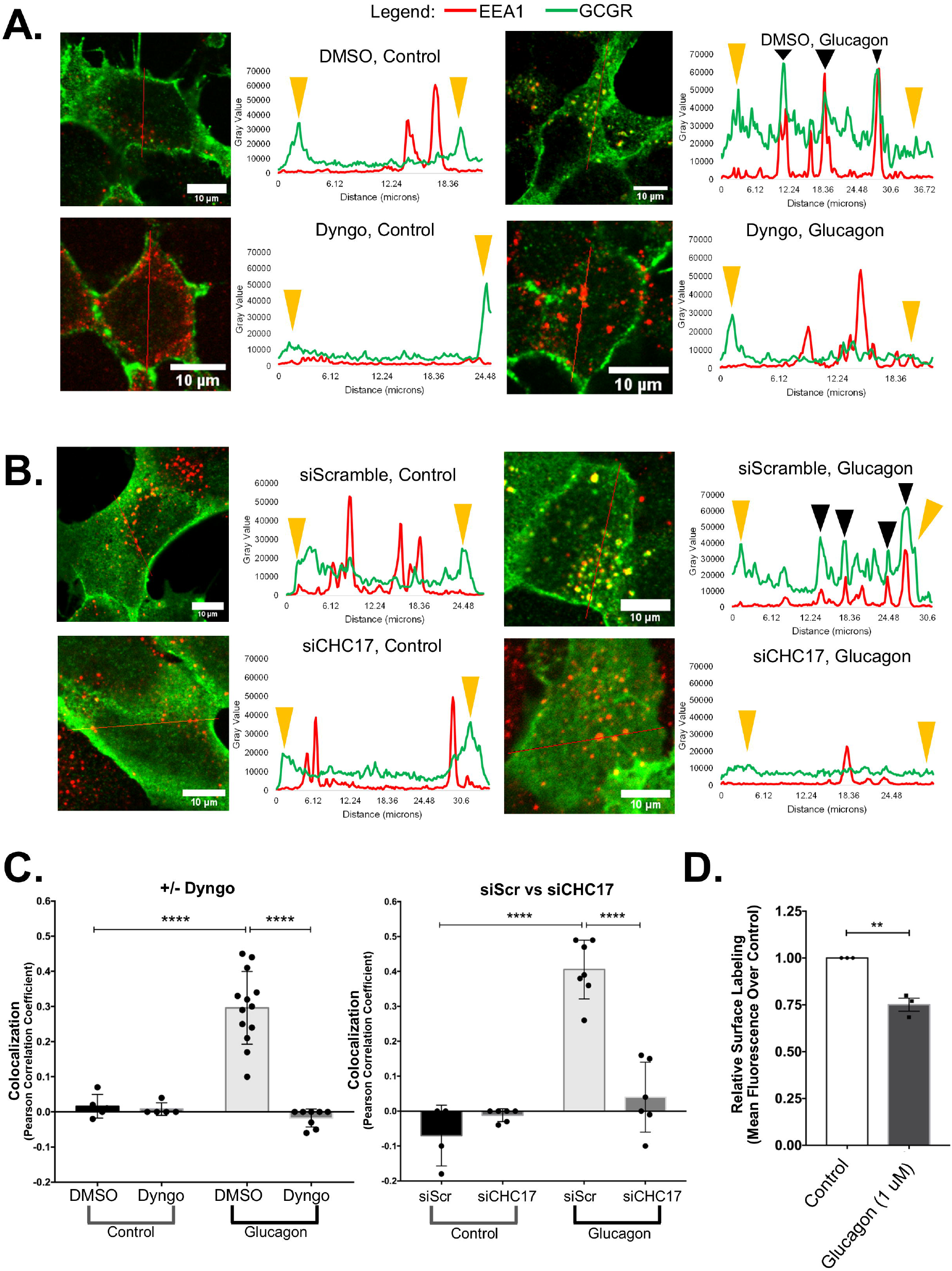
GCGR internalizes and localizes to endosomes upon stimulation with glucagon. HEK293 cells transiently transfected with GCGR-MycDDK were treated with or without 1 μM Glucagon for 30 minutes and analyzed by microscopy or flow cytometry. **(A-B)** Line scan analysis was performed on confocal microscopy images of cells labeled for GCGR (Green) and EEA1 (Red). Endocytosis was inhibited by either **(A)** pre-treatment for 15 minutes with or without 30 uM Dyngo or **(B)** transfection with siCHC17, or with siScramble as a control. GCGR localizes to the plasma membrane (yellow arrowheads), but translocates to endosomes upon stimulation (black arrowheads). Endocytic blockade via drug or siRNA prevented GCGR accumulation in endosomes. **(C)** Colocalization of GCGR and EEA1. Pixel intensities from confocal microscopy images were analyzed with the ImageJ Coloc 2 plugin. Left: Cells were pre-treated for 15 minutes with or without 30 uM Dyngo to block endocytosis. Right: Cells were transfected with siCHC17, or with siScramble as a control. GCGR co-localized with EEA1 after stimulation unless endocytosis was inhibited. All data are mean ± SD. Significance determined by ordinary one-way ANOVA followed by Tukey’s multiple comparisons test (**** p-value < 0.0001) **(D)** Upon 30 minute treatment with 1 μM glucagon, HEK293 cells stably expressing SSF-GCGR were labeled with M1 Flag antibody conjugated to Alexa Fluor 647, fixed and prepared for flow cytometry. Surface labeling of GCGR was measured. Mean fluorescence from 3 independent experiments were normalized to unstimulated control cells. All data are mean ± SEM. Significance determined by unpaired two-tailed t-test. (** p-value < 0.05)

GCGR and endosomal localization was further analyzed by calculating Pearson’s Correlation Coefficients (PCC) for GCGR and EEA1 (Figure 1C). GCGR colocalized with the endosomal marker EEA1 upon stimulation with glucagon. Inhibition of endocytosis, via treatment with 30 μM Dyngo, prevented GCGR from trafficking to the endosomes after glucagon stimulation. No significant differences were found by PCC when comparing unstimulated control cells with glucagon-stimulated samples with either Dyngo or siCHC17 treatment.

Finally, we performed surface-labeling assays to confirm internalization of GCGR. For these experiments, we treated HEK293 cells stably expressing SSF-GCGR with or without glucagon for 30 min. We then labeled the plasma membrane-localized GCGR using the M1 Flag antibody followed by FACS analysis (Figure 1D). Indeed, less surface-labeled GCGR was detected in cells treated with glucagon, as measured by a significant reduction in mean GCGR fluorescence intensity. This confirms that GCGR internalizes upon stimulation. Together, line scan, colocalization, and FACS analyses indicate that GCGR, upon stimulation with glucagon, internalizes and traffics via endocytosis.

### Endocytosis modulates GCGR Signal Transduction

After determining that GCGR traffics via endosomes in response to stimulation with glucagon, we sought to determine the requirement of endocytosis on GCGR-dependent signaling and downstream transcriptional regulation. Because HEK293 cells do not endogenously express GCGR, we generated stable cell lines that express the Flag-tagged receptor. Several GPCRs depend on endocytosis to achieve maximal transcriptional regulation, including the Beta 2 Adrenergic Receptor (B2AR) (Irannejad et al., 2013). In previous work, it was determined that B2AR, an endogenously expressed GPCR in HEK293 cells, signals in an endocytosis-dependent manner (Tsvetanova & von Zastrow, 2014). Additionally, the gluconeogenic enzyme Phosphoenolpyruvate Carboxykinase 1 (PCK1) was identified as an endocytosis-dependent transcriptional target of B2AR. For this study, we hypothesized that this gluconeogenic target is also regulated by endocytosis-dependent GCGR-signaling.Both GCGR and B2AR signal via the Gs subunit, and generate cAMP as a second messenger. Therefore, we included B2AR stimulation in our experiments as an internal positive control to verify that cAMP is produced in response to agonist stimulation, and that endocytic blockade inhibits Gs-mediated cAMP production.

To investigate whether GCGR signaling is endocytosis-dependent, we measured the production of cAMP, and the transcriptional regulation of *PCK1* in the presence or absence of endocytosis inhibition with Dyngo. Using a pGLO-based cAMP biosensor, we measured cAMP production over time in cells after treatment with glucagon or isoproterenol, in the presence or absence of Dyngo pre-treatment. Both isoproterenol and glucagon stimulated cells resulted in prominent cAMP production compared to untreated cells (Figure 2A). For both isoproterenol and glucagon-stimulated cells, relative luminescence over time was reduced in cells that were pretreated with Dyngo. Further analysis revealed that peak cAMP intensities in glucagon and isoproterenol-stimulated cells were significantly higher than cells pre-treated with Dyngo (Figure 2B). As previously published for B2AR stimulated cells, *PCK1* transcription was significantly induced by glucagon stimulation, unless cells were pretreated with Dyngo to inhibit endocytosis (Figure 2C). Together, these data indicate that the generation of maximal cAMP and the transcription of *PCK1* are dependent on the endocytosis of GCGR.

**Figure 2:**
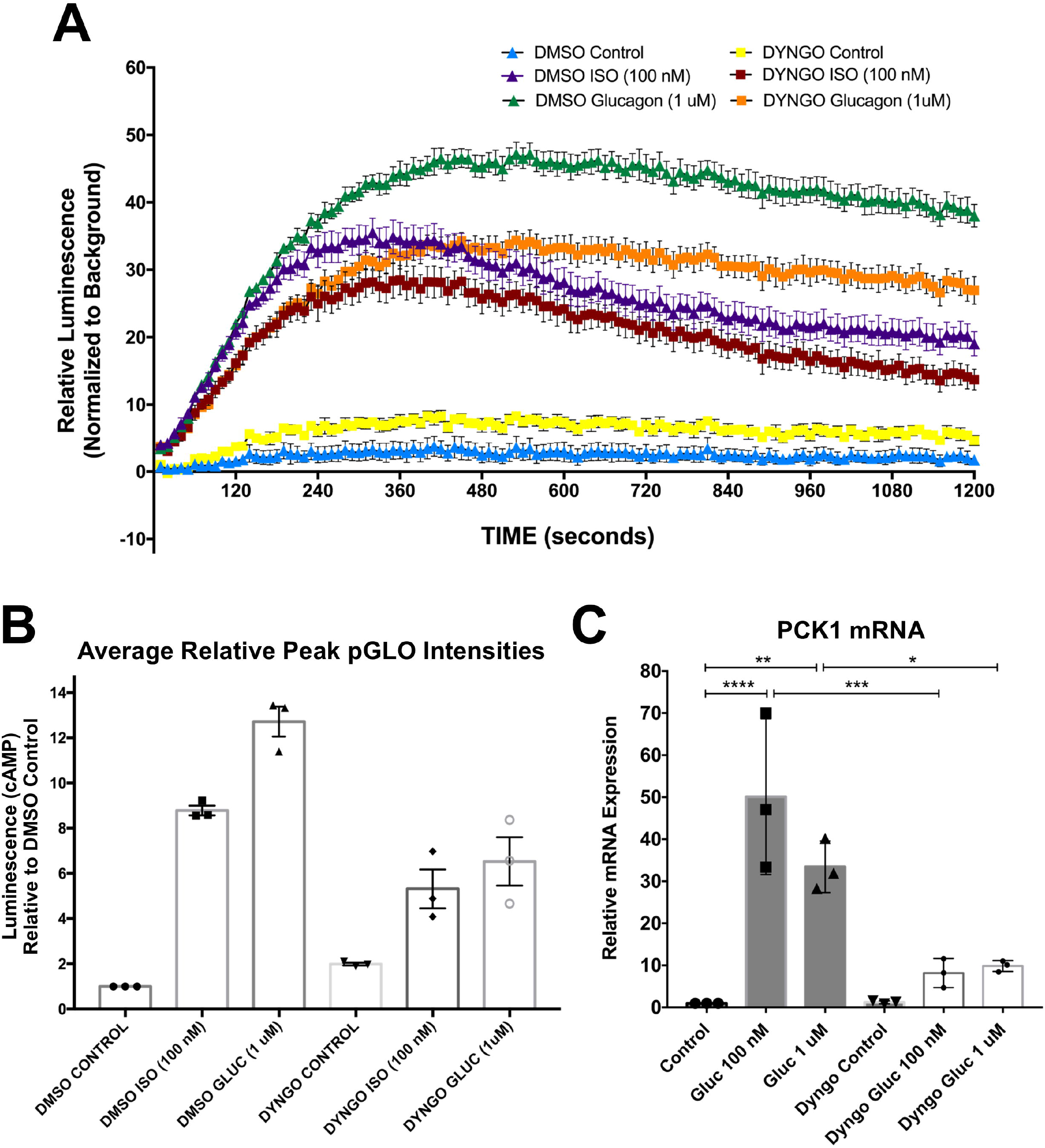
Maximal GCGR Signaling in Endocytosis Dependent. **(A)** HEK293 cells stably expressing SSF-GCGR were transfected with pGLO cAMP Biosensor. Cells were then pre-treated with or without 30 μM Dyngo. Upon treatment with or without 100 nM Isoproterenol or 1 μM glucagon, luminescence was measured every 10 seconds for 20 minutes. Representative data from one of three biological replicate experiments is shown. **(B)** Average peak intensities normalized to DMSO Control luminescence. **(C)** HEK293 cells stably expressing SSF-GCGR were pre-treated with or without 30 μM Dyngo to block endocytosis. Cells were then treated with or without 100 nM or 1 μM glucagon. RNA was isolated and qRT-PCR was used to quantify relative mRNA levels of the gluconeogenic gene *PCK1*. Endocytic blockade inhibits GCGR-dependent *PCK1* transcription. All data are mean ± SEM. Significance determined by ordinary one-way ANOVA followed by Tukey’s multiple comparisons test (**** p-value < 0.0001).

### Endocytosis is required for maximal GCGR-dependent transcriptional regulation in primary mouse hepatocytes

GCGR is endogenously expressed in the liver and serves as a regulator of blood glucose by upregulating gluconeogenic genes during hypoglycemic conditions (Janah et al., 2019). We sought to confirm that GCGR requires endocytosis for maximum transcription of gluconeogenic genes in a biologically relevant cell type. For this, experiments were conducted using primary mouse hepatocytes obtained from the UCSF Liver Center. Immediately following harvest, hepatocytes were seeded onto collagen coated cell culture plates. Twenty-four hours after plating, cells were pretreated with or without Dyngo and then treated with 100 nM glucagon for 2 hours. RNA was harvested for gene expression analysis by qRT-PCR. Gene expression was determined for the gluconeogenic genes, Phosphoenolpyruvate carboxykinase 1 (*PCK1*), Glucose-6-phosphatase (*G6PC*), and Peroxisome proliferator-activated receptor-gamma coactivator 1 alpha (*PGC1α*), in response to glucagon stimulation and inhibition of endocytosis. Gene expression of all three gluconeogenic targets was highly induced by glucagon stimulation, however, this upregulation was attenuated upon endocytic inhibition (Figure 3). Our data suggest that the transcription of gluconeogenic genes is endocytosis-dependent in primary hepatocytes.

**Figure 3:**
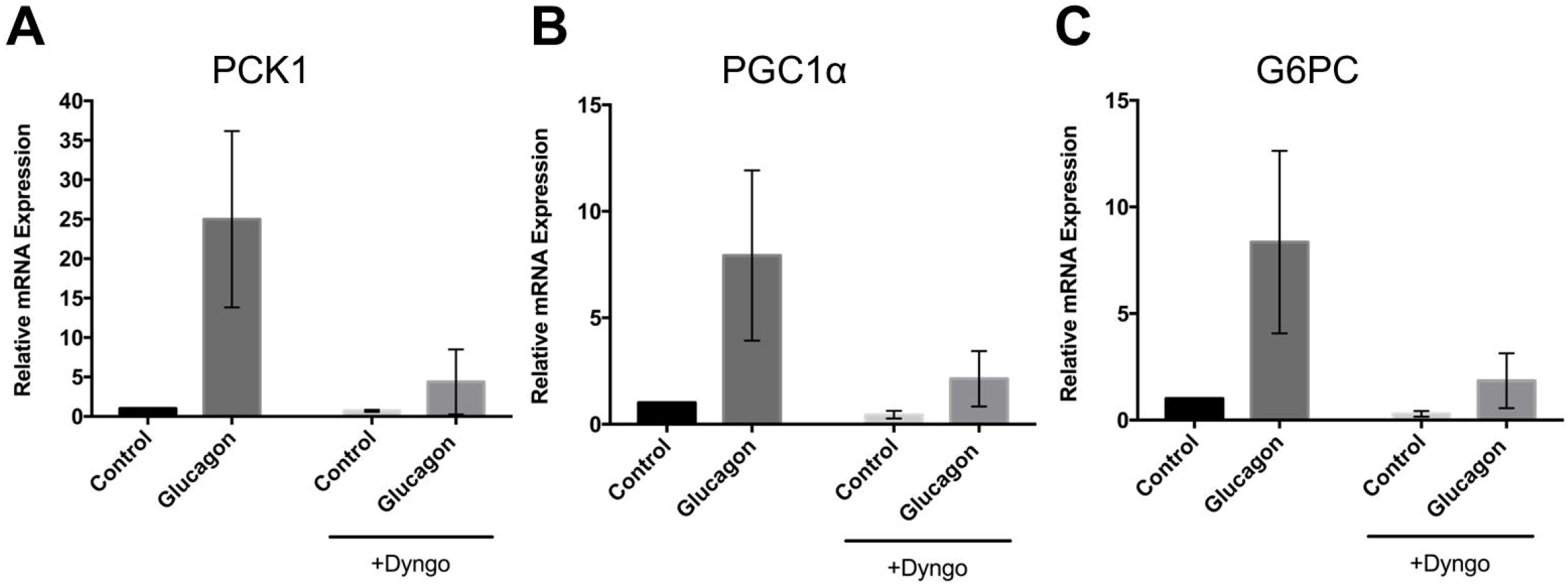
Endocytosis is required for maximal GCGR-dependent gluconeogenic gene transcription in primary mouse hepatocytes. Primary mouse hepatocytes were pre-treated or with or without 30 μM Dyngo for 15 minutes to block endocytosis before subsequent treatment with, or without 100 nM of glucagon for 2 hours. Relative mRNA expression was measured by qRT-PCR for gluconeogenic genes **(A)** *PCK1*, **(B)** *PGC1α*, and **(C)** *G6PC*.

## Discussion

Several studies have found that G protein-coupled receptor signaling after endocytosis is critical for maximal downstream regulation of transcription (Irannejad et al., 2013; Pavlos & Friedman, 2017; Tsvetanova et al., 2015; Tsvetanova & von Zastrow, 2014). The findings from our study demonstrate that upon activation by glucagon, GCGR must internalize for maximum transcription of gluconeogenic genes (Figure 4).

**Figure 4:**
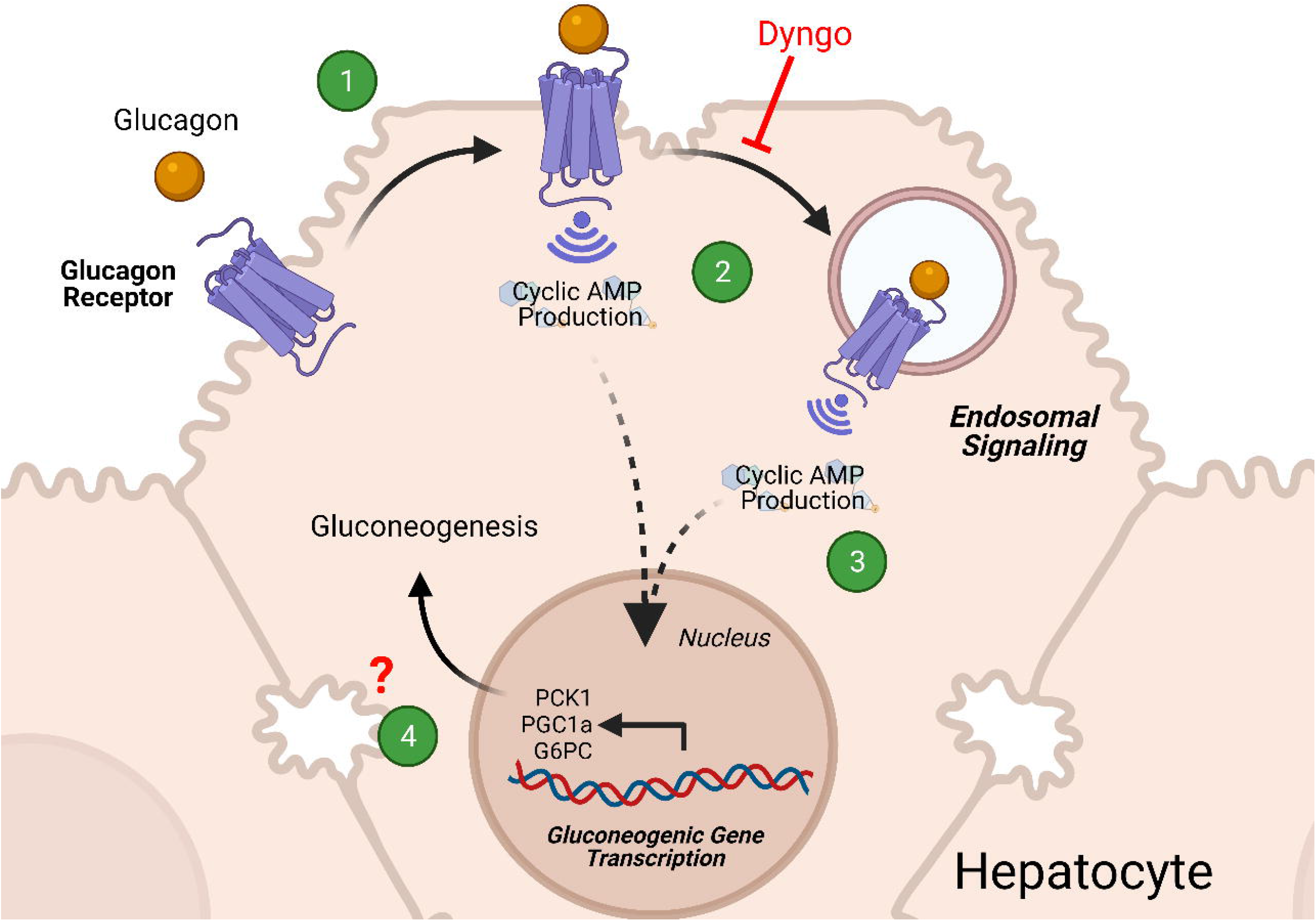
Model of endosomal Glucagon Receptor signaling and gluconeogenic gene regulation. **(1)** The peptide hormone glucagon binds to the GCGR, which is expressed in the plasma membrane of hepatocytes. This initial binding triggers cAMP generation at the plasma membrane. **(2)** GCGR is internalized via endocytosis. **(3)** Our data show that after endocytosis, GCGR also generates a significant amount of cAMP. **(4)** Endocytic-GCGR signaling is required for maximal transcription of gluconeogenic genes in hepatocytes.

First, we determined that upon ligand-dependent internalization, GCGR localizes with endosomes in HEK293 cells (Figure 1). Endocytosis is required for maximum GCGR signaling as measured by cAMP production and upregulation of *PCK1* in HEK293 cells (Figure 2). We then confirmed the importance of endocytosis in primary mouse hepatocytes, where we found that endocytosis is required for glucagon-dependent upregulation of *PCK1, PGC1α*, and *G6PC* (Figure 3). Together, our results support the notion that upon stimulation with glucagon, GCGR internalizes and localizes to the endosomes, and that this localization is required to exert maximal upregulation of gluconeogenic genes (Figure 4).

While we show localization to endosomes in HEK293 cells expressing GCGR treated with glucagon, previous studies have demonstrated that GCGR localizes to the endosomes in liver cells upon glucagon stimulation (Authier et al., 1992; Merlen et al., 2006). Other studies have explored GCGR internalization mechanisms. Previous work found that internalization in hepatocytes may not be mediated by Beta Arrestin (Krilov et al., 2008; Merlen et al., 2006). Another study done in CHO-K1 cells expressing exogenous Receptor Activity Modifying Protein 2 (RAMP-2) showed that GCGR internalizes with RAMP-2 instead of Beta-Arrestin, and that RAMP-2 increases GCGR-dependent cAMP generation (Cegla et al., 2017). Future experiments could confirm the requirement of RAMP-2 or Beta Arrestin by inhibiting either mechanism and interrogating gluconeogenic gene transcription after glucagon stimulation.

Subsequent studies should also explore the signaling activity of GCGR at the endosomal membrane, possibly using conformational biosensors, called nanobodies, specifically designed to bind activated GCGR. Nanobodies have been used to directly show that endosome-localized B2AR is in an active conformation (Irannejad et al., 2013). Moreover, GCGR may localize to additional intracellular compartments other than the endosomes. Therefore, we cannot rule out the possibility of cAMP generation from other membranes, as reported for the Thyroid Stimulating Hormone Receptor (Godbole et al., 2017). Determining specific mechanisms of subcellular GCGR signaling could inform future studies that explore new drug candidates in treating metabolic dysregulation.

Our findings expand on previous *in vivo* findings which showed a significant downregulation of gluconeogenic genes in mouse liver 5 days after the endocytic pathway was disrupted (Zeigerer et al., 2015). We focused on glucagon-specific signaling and found that inhibition of endocytosis has a notable impact on gluconeogenic gene transcription within minutes of treatment, thus refining our understanding of the implications of receptor endocytosis. Our work is the first to show that endocytosis of the Glucagon Receptor is required for maximal gluconeogenic gene transcription.

Glucagon and GCGR play important roles in blood glucose homeostasis and are an attractive target for therapeutic intervention in metabolism (Bagger et al., 2011). Despite this, the specific mechanisms of GCGR signaling have yet to be fully explored, particularly in the nuances of its subcellular signaling. These results highlight a key step between receptor activation and gluconeogenesis, and support the biological relevance of subcellular GCGR signaling. Together, our findings demonstrate an important molecular step in the physiological response to hypoglycemia.

## Acknowledgements

These studies were supported by the US National Institutes of Health (DA010711 and DA012864 to MvZ) and the UCSF Discovery Fund. ELS was supported by the NIH IRACDA (K12 GM081266/GM/NIGMS at UCSF) and the CSU Program for Education & Research in Biotechnology (CSUPERB) New Investigator Award at SFSU. JMC was supported by the San Francisco State NIH MBRS-RISE: R25-GM059298. We also thank Nina Tsvetanova, Grace Peng and the members of the von Zastrow lab for valuable advice and discussion. Finally, we would like to thank DeLaine Larsen, and Kari Herrington (Nikon Imaging Center, UCSF, National Institute of Health 1S10OD017993-01A1) for technical support and expertise, as well as, Jaquelyn J. Maher and Chris L. Her (UCSF Liver Center) for technical support and expertise in isolating primary mouse hepatocytes for these studies.

